# Epithelial stem cells in the premenopausal human vagina

**DOI:** 10.64898/2026.06.03.729864

**Authors:** Julia L. Balough, Kjiana E. Schwab, Temara M. Stransky, Tianjiao Chu, Abigail R. Cameron, Srividya P. Babu, Shanti Gurung, Krystyna Rytel, Caroline E. Gargett, Kyle E. Orwig, Pamela A. Moalli

## Abstract

The vagina undergoes physiologic changes across the menstrual cycle, pregnancy, birth and menopause. Most women will experience vaginal dysfunction at some point during their lives and treatment options are limited. The cyclic regeneration of the vaginal epithelium during each menstrual cycle suggests it is a stem-cell based tissue; however, human vaginal epithelial stem cells (veSCs) have not been identified. We utilized *in vitro* colony forming and organoid assays to confirm that cells in the human vaginal epithelium have self-renewal and differentiation potential. Specifically, we determined that stem cell activity resides in the ITGA6+ and NGFR+ fractions of the basal epithelium. We performed single-cell RNA sequencing to identify the distinct cellular compartments of the full thickness human vagina, including spatially distinct populations comprising layers of the stratified vaginal epithelium. Markers of these cells within the vaginal epithelium were validated by immunohistochemistry. CD9+ITGA6+NGFR+ and CD9+ITGA6+NGFR- cells were capable of efficient colony formation but only the CD9+ITGA6+NGFR+ fraction produced organoids containing basal, intermediate and superficial layers of the vaginal epithelium. We characterized the premenopausal human vagina at single cell resolution, validated markers and assays to test veSC developmental potential, and identified putative stem cells that may open new avenues for treating vaginal dysfunction.

## Introduction

The vagina is a dynamic organ composed of epithelial, mesenchymal and connective tissues that undergoes remodeling across the lifespan of women. Vaginal remodeling occurs in response to hormonal changes associated with puberty, the menstrual cycle, pregnancy, and after birth. These are necessary to facilitate pH balance, a *Lactobacillus*-dominant microbiome, protection from pathogens, blood flow, elasticity and lubrication, all of which are necessary for healthy vaginal function^1^. Vaginal dysfunction can arise from congenital sources such as Mayer-Rokitansky-Kuster-Hauser (MRKH) syndrome where approximately 1 in 5000 XX individuals are born with partial or complete absence of vagina^2^. Women who undergo cancer-related procedures such as hysterectomy for cervical^3^ or endometrial cancer^4^ may have portions of the vagina removed. In addition, treatments such as radiation or chemotherapy can confer indirect impacts, arising from ovarian dysfunction, and direct impacts on vaginal function^5^. Finally, all women will undergo menopause, of which 70% will seek treatment for the genitourinary syndrome of menopause (GSM)^6^. GSM is a naturally occurring, chronic condition caused by the loss of estrogen production by the ovary with age. Individuals may experience vaginal dryness, burning, irritation, dyspareunia, dysuria, and lower urinary tract dysfunction which are the result of vaginal epithelium thinning, loss of epithelial glycogen, reduced vascularity, and an altered extracellular matrix composition^7,8^. Beyond symptomatic manifestations, vaginal dysfunction at any stage of life impairs sexual and urinary function, psychosocial well-being, and quality of life^9,10^.

The most common treatment for vaginal conditions includes local estrogen therapy delivered via creams, tablets, or rings and can be effective in restoring epithelial maturation and alleviating symptoms^11^. However, patient adherence is low with discontinuation rates exceeding 50% within the first year of use, often due to concerns about hormone exposure, inconvenience, cost, or social stigma surrounding vaginal atrophy^12,13^. Moreover, estrogen therapy does not alleviate symptoms or is not appropriate for all women, particularly women who have received pelvic irradiation, systemic chemotherapy, or endocrine treatments^5,14,15^. These limitations underscore the need for hormone-free, biologically driven approaches to restoring vaginal health.

Tissue-resident stem cells (SCs) may offer a regenerative medicine intervention for vaginal atrophy. Stem-cell based therapies are emerging in gynecologic medicine^16^; however, these therapies often use stem cells from non-autologous or non-endogenous tissue sources such as mesenchymal stem cells derived from bone marrow^17^ or umbilical cord^18^ for vaginal repair due to pelvic organ prolapse or vulvar tissue for vaginal aplasia^19^. While these approaches do show benefit in tissue repair after injury, this may be attributed to the anti-inflammatory properties of MSCs rather than resident SC-based regeneration of a healthy vaginal tissue^20^. There is a need to identify and characterize endogenous vaginal stem cells that can be therapeutically harnessed for gynecologic regenerative medicine applications.

The vagina, a hormonally responsive organ, undergoes cyclical epithelial remodeling during each menstrual cycle throughout the reproductive years and self-repairs after vaginal birth injury, suggesting it is a stem cell-based epithelium. A population of putative stem cells marked by CD271 (NGFR) and *Axin2* were discovered in the mouse vaginal epithelium that are capable of self-renewal in the absence of estrogen and differentiation upon estrogen re-exposure^21^. To date, human vaginal epithelial stem cells have not been identified. Published transcriptional profiles of the human vagina are derived from fetal^22,23^, post-menopausal or patients with pelvic organ prolapse^24^; thus, it is necessary to establish a comprehensive transcriptional and cellular map of the premenopausal human vagina. We hypothesize the human vagina contains tissue-resident epithelial stem cells that self-renew and differentiate to maintain a functional vaginal epithelium or regenerate the epithelium after trauma.

To enable studies of the human vagina and specifically vaginal epithelial stem cells (veSCs), we established assays to test self-renewal by colony formation and differentiation in epithelial organoids, *in vitro*. We generated a transcriptomic atlas from single cell RNA sequencing of full-thickness, pre-menopausal, non-prolapse human vaginas. We utilized this map to define distinct cell populations in the human vagina and predict a developmental trajectory from stem cells to terminally differentiated cells in the vaginal epithelium. Further, we used the single cell RNAseq data to establish a catalog of markers specific to each spatially distinct cell population or cellular compartment of the human vaginal epithelium, which were then validated by immunohistochemistry. The cell surface markers, integrin-alpha 6, ITGA6 (CD49f), and nerve growth factor receptor, NGFR (CD271), were expressed by cells on the basement membrane of the vaginal epithelium and were used to isolate subpopulations of basal epithelial cells. The *in vitro* stem cell assays confirmed that the human vaginal epithelium contains cells with the hallmark stem cell characteristics of self-renewal and differentiation. Stem cell activity was limited to a specific ITGA6+NGFR+ subpopulation of human vaginal epithelial cells.

In summary, we established a transcriptional atlas of the premenopausal adult human vagina and validated experimental tools to enable future studies of the human vagina in normal or disease states. The human vaginal epithelium is a stem cell-based tissue, suggesting the possibility of regenerative medicine approaches to treat vaginal dysfunction, which will impact all women at some point in their lives.

## Materials and Methods

### Human Tissue

Full thickness biopsies of the vagina were obtained from premenopausal, ages 25-52, women undergoing benign gynecologic surgery under IRB-approved protocols at the University of Pittsburgh (#19080175, n=10 samples) or Monash Health (#78468, n=25 samples) (Table 1). Written informed consent was obtained prior to surgery for each participant. Eligibility criteria included women with ovaries, who reported regular cycling over the previous 12 months, undergoing pelvic procedures such as tubal ligation, dilation and curettage, hysterectomy, vaginal wall repair, mid-urethral sling procedures, cystoscopies, peri-urethral bulking, or pelvic floor muscle injections for chronic pelvic pain. Vaginal tissues were acquired intraoperatively either using a 12 mm punch biopsy or undermining and excising with Metzenbaum scissors. Vaginal biopsies collected from the University of Pittsburgh Medical Center Magee-Womens Hospital (UPMC) were placed into dPBS (ThermoFisher, 14190144) and transported on ice to the laboratory for immediate processing. Vaginal tissue collected from Monash Health hospitals were placed in collection media comprising DMEM/F12 HEPES media (Gibco, 11330-057), 5% newborn calf serum (NCS, Gibco, 16010-159) and 1% antibiotic-antimycotic (anti-anti, Gibco, 15240-062), stored at 4°C and processed within 24 hours. Tissue samples were allocated for clonogenicity assays (n=26) and/or organoid formation (n=18) or single cell RNA sequencing (n=6). For larger specimens, a portion of the sample was kept intact and fixed for marker validation as described below.

**Table 1.** Sample Age, Surgical Indication and Assay Allocation.

|  | Age<br>(Years) | Surgical Indication | Assay |  |  |
| --- | --- | --- | --- | --- | --- |
|  |  |  | scRNAseq* | CFU | Organoid |
| Human<br>Samples<br>N=35 | 25 | Endometriosis | X (A) |  |  |
|  | 30 | Menorrhagia | X (B) |  |  |
|  | 32 | Urinary Incontinence |  | X | X |
|  | 32 | Menorrhagia, Sterilization |  | X |  |
|  | 33 | Sterilization | X (F) |  |  |
|  | 34 | Menorrhagia, Pain |  | X | X |
|  | 35 | Pelvic Organ Prolapse |  | X |  |
|  | 36 | Pelvic Organ Prolapse |  | X | X |
|  | 38 | Pelvic Organ Prolapse |  | X | X |
|  | 39 | Sterilization | X (C) |  |  |
|  | 39 | Pelvic Organ Prolapse |  |  | X |
|  | 40 | Menorrhagia, Sterilization |  | X | X |
|  | 40 | Uterine Fibroid, Cyst, Vaginal Septum |  | X | X |
|  | 40 | Pelvic Organ Prolapse, Urinary Incontinence |  | X | X |
|  | 40 | Pelvic Organ Prolapse, Menorrhagia |  | X | X |
|  | 41 | Pain | X (E) |  |  |
|  | 41 | Vaginal Repair |  | X |  |
|  | 41 | Pelvic Organ Prolapse, Urinary Incontinence |  | X | X |
|  | 41 | Pelvic Organ Prolapse |  | X | X |
|  | 41 | Pelvic Organ Prolapse |  |  | X |
|  | 42 | Pelvic Organ Prolapse |  | X |  |
|  | 43 | Uterine Fibroid, Pain | X (D) |  |  |
|  | 43 | Uterine Fibroid, Pain |  | X |  |
|  | 43 | Pelvic Organ Prolapse |  |  | X |
|  | 44 | Pelvic Organ Prolapse, Menorrhagia, Pain |  | X |  |
|  | 45 | Menorrhagia, Endometriosis |  | X | X |
|  | 45 | Pelvic Organ Prolapse |  | X |  |
|  | 45 | Pelvic Organ Prolapse, Urinary Incontinence |  | X |  |
|  | 46 | Menorrhagia, Uterine Fibroid, Pain |  | X |  |
|  | 47 | Menorrhagia |  | X |  |
|  | 47 | Pelvic Organ Prolapse |  | X | X |
|  | 48 | Urinary Incontinence |  | X |  |
|  | 50 | Pelvic Organ Prolapse, Urinary Incontinence |  | X | X |
|  | 51 | Pelvic Organ Prolapse |  | X | X |
|  | 52 | Pelvic Organ Prolapse |  | X | X |
| Median | 41 |  | N=6 | N=25 | N=18 |

\*Letter indicates the sample UMAP in Supp. Fig. 1

### Tissue Digestion

Vaginal tissue was washed in sterile dPBS (ThermoFisher, 14190144) then placed in a pre-weighed tube of collection media and weighed again. Tissue was washed with sterile dPBS and cut into 5 x 7 mm segments and placed into dPBS with 4 mg/mL Dispase II (Thermo Fisher, 17105041) and 500 mM Glucose (MP Biomedicals, 194024) at a ratio of 5 mL of solution for 500 mg of tissue. Tubes containing tissue were agitated (3 x g) on a shaker at 37°C for 60-90 minutes. Pieces of vagina were placed in a petri dish and the epithelium was gently peeled off the fibromuscular mesenchyme with forceps and each portion was placed into fresh dPBS, finely minced, then centrifuged (600 x g) for 5 minutes. Minced epithelium was then transferred to 10 mL of pre-warmed TrypLE Express (ThermoFisher, 12604013) and agitated at 37°C for 10-15 minutes, before any remaining clumped epithelium was broken down to single cells by mechanical means for 3-5 minutes. Meanwhile, minced mesenchyme was transferred to 15 mL of dPBS containing 20 mg/mL Collagenase I (Worthington, CLS-1), 4 mg/mL DNase I (Worthington, DP), and 500 mM Glucose and agitated at 37°C for 30-45 minutes. Every 15 minutes, the mesenchyme was triturated by pipetting up and down. When single cells were visualized in both digestions, the digestion was quenched by adding an equal part of collection media to each digestion solution. Mesenchymal cells were sequentially filtered through a 100 μm (Falcon, 352360) and 40 μm cell strainers (Fisher Scientific, 22-363-547). Epithelial cells were filtered through a 40 µm cell strainer only. Collected cells were centrifuged (600 x g) for 5 minutes. The supernatant was removed and the pellets were resuspended. Cell count and viability with trypan blue (Gibco, 15250-061) was performed using a hemocytometer. Cells were used fresh for *in vitro* assays, described below, or resuspended in Recovery™ Cell Culture Freezing Medium (Thermo Fisher, 12648010), frozen slowly overnight in a foam cooler at -80°C and transferred to liquid nitrogen for long term storage.

### Vaginal Fibroblast Feeder Layer

Mesenchymal cells isolated as described above were thawed and cultured in standard tissue culture flasks at 37°C and 5% CO_2_ in DMEM/F12 supplemented with 10% FBS and 1% anti-anti (fibroblast media)^25^. Cells were expanded through 2-3 passages and grown to confluence. Cells were then treated with 10 µM Mitomycin C (Stemcell Technologies, 73274) for 3-4 hours at 37°C in 5% CO_2_. After washing with dPBS, the cells were lifted with 0.05% Trypsin-EDTA (Thermo Fisher, 25300062) for 10 minutes. Trypsin was inactivated by the addition of an equal part fibroblast media, then the solution was centrifuged at 600 x g for 5 minutes and the cells pelleted. After aspiration of the supernatant, the pellet was resuspended for colony forming assays.

### Colony Forming Assay

Fibronectin coated dishes seeded with a mitotically-inactivated fibroblast feeder layer were prepared for the colony forming assay 1-3 days in advance. 100 mm petri dishes (Corning, 430167) were coated with 5 mL of fibronectin (Thermo Fisher, 33016015) in dPBS at 37°C for at least 20 minutes. After aspirating the excess nonadherent fibronectin solution, Mitomycin C treated feeder cells suspended in fibroblast media were plated at a density of 4000 cells/cm^2^. After tissue digestion, cell count, and cell sorting (where applicable), vaginal epithelial cells were resuspended in epithelial cell media composed of OptiPRO Serum Free Media (ThermoFisher, 12309019) supplemented with 2.5% CTS KnockOut SR XenoFree Media (ThermoFisher, 12618012), 1% L-Glutamine (ThermoFisher, 25030081 or Gibco, 35050-038), 1% anti-anti and 1 µM of the TGF-β pathway inhibitor, A83-01 (Stemcell Technologies, 72022) and plated at a density of 1500 cells/100 mm plate and incubated at 37°C in 5% CO_2_/95% air. Media was changed every 5-6 days. Cultures were terminated after 14 days by fixation in 10% formalin (Fisher Scientific, SF98-4) for 10 minutes. Plates were washed in dPBS then stained with 5% Crystal Violet solution (Millipore Sigma, V5265) for 1 minute. Plates were washed with tap water 5 times and dried at RT. Colonies were defined as clusters of cells greater than ∼1mm in diameter. Plates were scanned with a Konica Minolta Bizhub 300i (Konica Minolta Business Solutions, New Jersey, USA). The number of colonies formed from single cells was counted macroscopically per plate and the efficiency (%) was determined by calculating the number of colonies formed divided by the number of cells seeded (1500) x 100.

### Organoid Formation

Epithelial cells (5000 cells/dome) were seeded in 30 µL of growth factor reduced Matrigel for Organoid Culture (Corning, 356255) onto culture-treated coverslips (Nunc, 174950) in a 24-well plate. Matrigel domes were allowed to polymerize at 37°C for 30 minutes and then overlayed with vaginal organoid media (VOM)^26^ containing Advanced-DMEM/F12 (Thermo Fisher, 12634010) supplemented with 12.5% noggin conditioned media (Millipore Sigma, SCM105 or produced by Monash Biomedicine), 2 µM A83-01 (Tocris, 2939), 1% anti-anti, 1% L-Glutamine, 1% B27 Supplement (Gibco, 17504044), 1% N2 Supplement (Gibco, 17502048), 0.5 µg/ml hydrocortisone (Sigma-Aldrich, H0888), 10 nM nicotinamide (Sigma-Aldrich, N0636), 1.25 mM N-acetyl-l-cysteine (Sigma-Aldrich, A9165), 10 µM forskolin (Sigma-Aldrich, 344282), 10 ng/ml human epidermal growth factor (EGF, Gibco, PHG0311L), 100 ng/ml human fibroblast growth factor-10 (Peprotech, 100-26) and 100 ng/ml Y-27632 ROCK pathway inhibitor (Sigma-Aldrich, Y0503). The ROCK inhibitor was removed from VOM at day 3 and media was changed every 3 days. Matrigel domes with epithelial cells were cultured for 10-12 days and were imaged at day 12 of culture with transmitted light at 4X and 20X.

### Organoid and Tissue Fixation and Processing

After 10-12 days in culture, organoids were fixed for OCT (Fig. 1) or FFPE (Fig. 6). For OCT embedding, Matrigel domes were fixed in zinc formalin (Sigma-Aldrich, Z2902) for 5 minutes, then immersed in O.C.T. compound (Sakura, 4583), before being snap frozen on dry ice and stored at -80°C. Intact vagina was fixed in 4% Paraformaldehyde (Sigma-Aldrich, P6148) overnight at 4°C, then transferred to 30% sucrose in PBS (Sigma-Aldrich, S0389) for 48 hours and embedded in O.C.T compound and stored at -80°C for future analysis. Cryosectioning of OCT embedded veOrganoids and vaginal tissue and hematoxylin and eosin staining were performed by the Monash Histology Platform (MHP).

**Figure 1.**
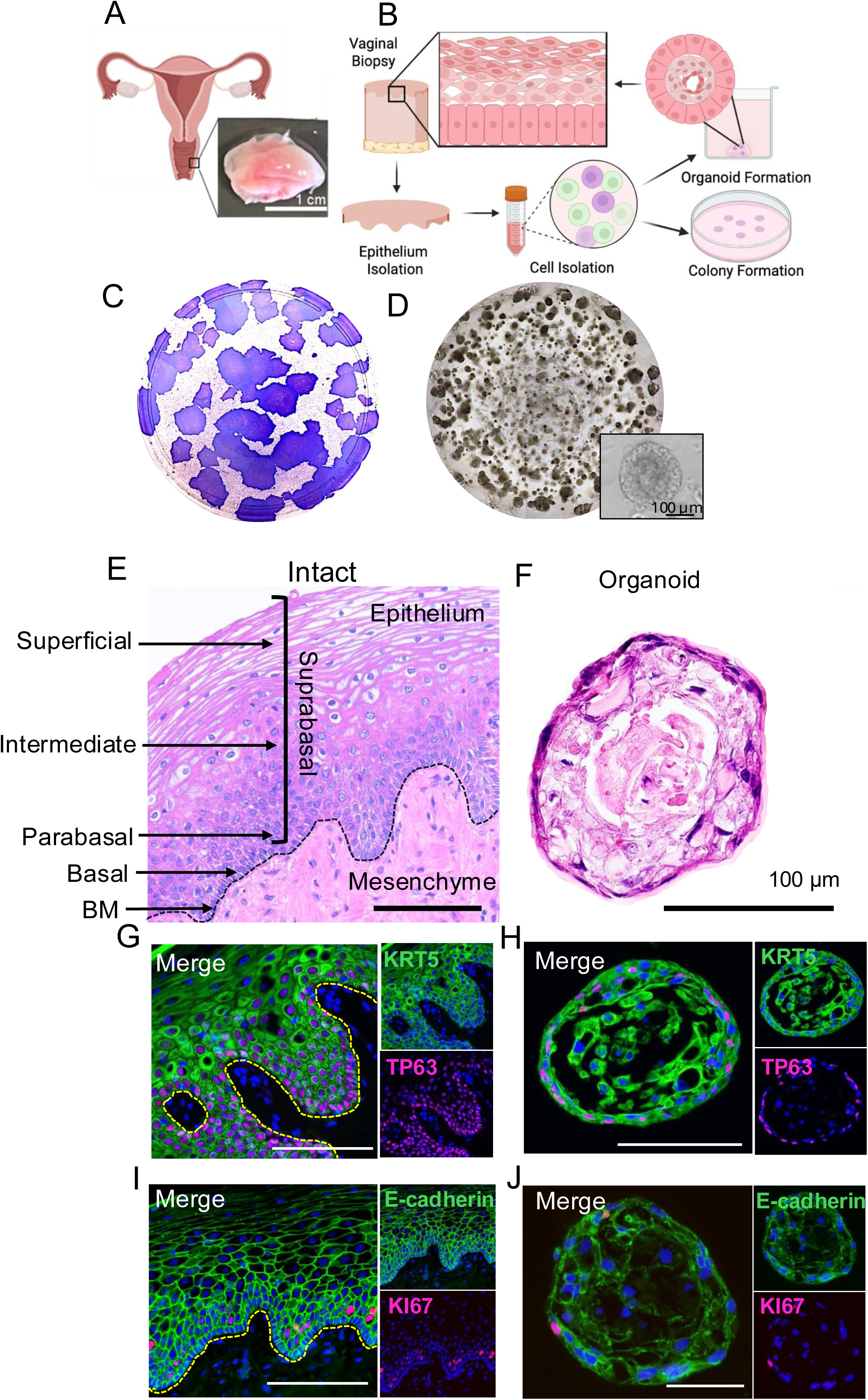
From single cells, the human premenopausal vaginal epithelium can form colonies and create organoids that resemble the intact epithelium. A) Representative full thickness biopsy (∼1 cm^2^) from the premenopausal human vagina. B) Experimental schematic for isolation of vaginal epithelial cells used in colony or organoid formation assays. C) Representative Crystal Violet stained culture plate displaying colonies formed 14 days after single cells were seeded (1500 cells/plate). D) Representative transmitted light image of Matrigel dome containing vaginal epithelial organoids formed after 10-12 days in culture. E) Histologic description of the hematoxylin and eosin stained intact premenopausal human vaginal epithelium. The epithelium is a stratified squamous epithelium on a mesenchymal layer. The epithelium contains basal cells adhered to a basement membrane (BM) and suprabasal cells which contain parabasal cells, intermediate cells, and superficial epithelial cells. F) Representative vaginal epithelial organoid, derived from unsorted vaginal epithelial cells from the premenopausal vagina, stained with hematoxylin and eosin. G) The intact vaginal epithelium displays multiple cell layers stained with KRT5 (green) and basal cells with TP63 (magenta) adjacent to a basement membrane (dashed yellow line). H) veOrganoid displays KRT5 (green) epithelial cells throughout the organoid with TP63 (magenta) around the periphery. I) The intact vaginal epithelial displays E-cadherin (green) throughout the basal and suprabasal layers with sporadic KI67 (magenta) cells primarily in the parabasal region. J) veOrganoids display E-cadherin (green) throughout the organoid with KI67 (magenta) cells on the exterior.

For FFPE, Matrigel domes were fixed in Modified Davidson solution (Electron Microscopy Sciences, 64133-50) for 4 hours at RT, then overnight at 4°C. Matrigel domes were washed with 70% ethanol and carefully transferred into cassettes. Processing and embedding of veOrganoids in Pittsburgh was performed by the Histology and Microimaging Core at Magee-Womens Research Institute. In brief, Matrigel domes were wrapped in lens paper and sandwiched between two sponges within the cassette for processing. Tissue processing was performed using a Tissue-Tek VIP Tissue Processor (Sakura FineTek USA, Torrance, CA, USA). Domes were paraffin embedded and sectioned into 5 µm thickness. Organoids were stained with hematoxylin and eosin (H&E) to visualize architecture by the Pitt Biospecimen Core (UPMC Shadyside, Pittsburgh, PA, USA). Intact vagina was fixed, processed and embedded following the same protocol for histologic analysis and marker validation by immunohistochemistry.

### Immunohistochemistry

Immunohistochemistry was performed on intact premenopausal vagina to validate marker protein localization as well as organoids to determine the cell composition after culture.

For OCT blocks, defrosted slides were fixed in zinc formalin and then subjected to heat-induced epitope retrieval according to standard protocols by the MHP. Sections were permeabilized using 0.1% Tween/PBS (Sigma-Aldrich, P1754), washed twice in PBS, then incubated with serum-free protein blocking solution (Dako, X0909) for 30 min at RT. Primary antibody solutions were prepared in 1% Bovine Serum Albumin/PBS (BSA, Sigma-Aldrich, A2153). We labelled epithelial cells with CD9-FITC (mouse, BD Biosciences, 555371), E-cadherin (mouse, BD Biosciences, 610181) and cytokeratin 5 (rabbit, Abcam, ab52635). Proliferating cells were stained with Ki67 (rabbit, Invitrogen, MA5-14520) and P63 (mouse, Abcam, ab735). Putative epithelial stem cells were labelled with ITGA6 (rat, BD Biosciences, 555734) and NGFR (mouse, Invitrogen, 14-9400-82). Following the primary antibody incubation overnight at 4°C in a humidity chamber, slides were washed twice in PBS before incubation with fluorescently conjugated secondary antibodies (goat anti-mouse IgG AF488 and AF568 (Invitrogen, A11001 and A11004), donkey anti-rabbit IgG AF568 (Invitrogen, A10042) and goat anti-rat IgG BF488 (Bioss, bs-0293G-BF488) prepared in 1% BSA/PBS for 1 hour at RT. Next, slides were washed 3 times in PBS on a rotor, followed by 0.05% Hoechst nuclear stain in MilliQ water (Invitrogen, H3569) for 5 min at RT, washed twice in MilliQ water and mounted using fluorescence mounting media (Dako, S3023). Staining was visualized using an Olympus BX53 fluorescence microscope and CellSens software (Olympus) using 20X and 40X objectives.

For FFPE tissue, slides were warmed at 60°C for 10 minutes and deparaffinized in 3 washes of Xylene for 5 minutes each. Next slides were rehydrated in sequential ethanol baths (5 minutes each): 2 x 100% EtOH, 95% EtOH, 80% EtOH, 70% EtOH, 50% EtOH, 25% EtOH then washed in 1X PBS for 2 minutes. Slides underwent antigen retrieval in TRIS (100 mM Tris and H_2_O at pH 10.0) at 95.5°C for 45 minutes then removed and allowed to cool for 30 minutes before washing twice in 1X PBS (2 minutes each). Slides were permeabilized in 0.02% Triton X-100 with agitation for 30 minutes then washed PBS-T (0.1% Tween-20 and 1X PBS) for 5 minutes each. Tissue sections were blocked with Donkey Blocking Buffer (5% Normal Donkey Serum, 3% BSA, 0.1% Triton X-100, 1X PBS) for 60 minutes at RT in a humid chamber. Blocking buffer was removed and tissue sections were incubated with primary antibody (1:100) diluted in donkey blocking buffer overnight at 4°C. CD9 (rabbit, Abcam, ab307085) was used to mark epithelial cells. We utilized ITGA6 (rabbit, SigmaAldrich, HPA012696) and COL7A1 (mouse, MilliporeSigma, SAB4200686) for basement membrane visualization. For basal cells, we utilized: KRT19 (rabbit, Invitrogen, HPA002465), COL17A1 (mouse, MilliporeSigma, MABT1521), and CDH13 (goat, R & D Systems, AF3264). For proliferative cells, we used PCNA (rabbit, Abcam, ab92552) and KI67 (mouse, BD Biosciences, 550609). For differentiating cells, we used KRT1 (rabbit, SigmaAldrich, HPA017917), KRT4 (rabbit, SigmaAldrich, HPA034881) and TGM1 (rabbit, SigmaAldrich, HPA070372). After primary antibody incubation, slides were washed 3 times in PBS-T for 5 minutes each and incubated with secondary antibody (1:200 in Blocking Buffer) for 45 minutes at RT. Secondary antibodies (1:200) used were donkey anti-rabbit IgG Alexa Fluor 568, donkey anti-goat IgG Alexa Fluor 488 (Invitrogen, A32814), and donkey anti-mouse IgG Alexa Fluor 488 (Invitrogen A21202). Slides were washed 3 times in PBS-T for 5 minutes each, then once in 1X PBS for 5 minutes. Finally, slides were mounted in Vectashield PLUS with DAPI (Vector Laboratories, H-2000-10) and coverslipped. Images were taken using a Nikon Eclipse 90i upright fluorescent microscope using 20X and 40X objectives.

### Fluorescent-Activated Cell Sorting (FACS)

FACS of freshly isolated human vaginal epithelial cells (described above) were transported in dPBS with 5% FBS. First, 10% of the cell solution was separated as an unstained control and left on ice. The remaining cells were then stained with FITC-conjugated mouse anti-CD9 (BD Pharmingen 555371, 1:45 dilution), APC-conjugated rat anti-ITGA6 (Miltenyi 130-097-250, 1:90 dilution), and PE-conjugated mouse anti-NGFR (Miltenyi 130-113-421, 1:450 dilution) on ice for 25 min. After incubation, stained and unstained cells were centrifuged (600 x g) for 5 minutes, the supernatants aspirated, and the pellets resuspended in 300 µL FACS buffer. Then the cell solution was filtered through a 35 µm filter cap (Falcon, 352235) and kept on ice until sorted. SYTOX Blue Dead Cell Stain (Molecular Probes S34857, 1:300 dilution) was added 5 min prior to sorting.

Isotype controls were made using rat BD comp beads (BD Biosciences, 51-90-9001189) for APC and mouse BD comp beads (BD Biosciences, 51-90-9001229) for the remaining fluorochromes. The beads were vortexed briefly, then one drop of both Ig-k positive and negative beads was added to a 15 mL tube for each fluorochrome. Isotype control antibodies of the matching fluorochrome were added to each tube as follows: 20 µL FITC anti-mouse IgG (BD Biosciences, 555748), 10 µL APC anti-rat IgG (Invitrogen, 17-4321-81) or 2 µL PE anti-mouse IgG (BD Biosciences, 554680). The beads were then incubated, centrifuged, resuspended, and filtered identical to the epithelial cells as described above. Stained beads were kept on ice until sorting.

Cell sorting proceeded immediately after staining using a BD Biosciences FACS Aria III and BD FACSDiva software (version 8.0.1). Cells were selected using electronic gating according to their forward versus side scatter profile. Doublets were excluded using a side scatter height versus side scatter width profile, then a forward scatter height versus width profile. Dead cells were excluded based on SYTOX Blue fluorescence.

### Single-Cell Library Preparation and Sequencing

Single cell library preparation and sequencing was performed at the High Throughput Genomics Core and Single Cell Core Facility at the University of Pittsburgh. After tissue dissociation, cell suspension concentration and viability were checked using ViaStain Acridine orange and propidium iodide (AOPI) staining solution (Revvity, CS2-0106) on the Revvity Cellometer Ascend (Revvity Inc, Waltham, MA, USA). Libraries are then prepared according to the manufacturer’s protocol using Chromium GEM-X Single Cell 3’ Reagent Kits (10X Genomics). The resulting full-length cDNA was checked for size and concentration on the Agilent Tapestation 4150 (Agilent, Santa Clara, CA, USA) before proceeding with gene expression library preparation and sample indexing. Final libraries were checked for size on the Agilent Tapestation and concentration using the Invitrogen Qubit 2.0. (ThermoFisher). Libraries are then sequenced on a NovaSeq X Plus sequencer (Novogene Corporation Inc, Sacramento, CA, USA).

### Single-Cell RNA Sequencing Analysis

The FASTQ files from 10X Genomics were processed by Cell Ranger (version 6.1.1), using the CellRanger count pipeline. Specifically, after trimming and filtering, the single-cell RNA seq data were aligned to human reference genome GRCh38 using aligner STAR. Reads that aligned to exons and introns in the sense orientation were selected and processed to generate unique molecular identifier (UMI) counts for the genes. In total we obtained 129,527 cells with 36,601 genes expressed in one or more cells. The gene count data were further analyzed using R package Seurat (v4). We first filtered low quality cells, defined as cells with less than 500 genes, or less than 2500 total reads, or ≥ 20% of the reads from mitochondrial genes. In total 51,882 cells passed the quality filtering (Supp. Fig. 1). Note that two samples (E and F) have only about 10% of cells passing the quality filtering (Table 1, Supp. Fig. 1B). We further filtered low expression genes, defined as genes expressed in less than 1% of cells. In total 15,193 genes passed this filtering step. The filtered data were then normalized and log transformed. The top 2000 genes with excessive variance among the cells analyzed were identified. Principal component analysis (PCA) was performed on log transformed normalized expression of these 2000 genes. Using the elbow plot, we selected the top 12 significant principal components. Based on the selected significant principal components, the cells were clustered into 26 clusters using a graph-based clustering algorithm implemented in Seurat. UMAP plots were generated to visualize the cell clusters^27^. Wilcoxon rank sum tests were used to determine the genes differentially expressed between cells from one cluster and cells from all other clusters. Top genes up regulated in each cluster were chosen as marker genes for the cell type represented by the corresponding cluster. Bulk gene differential expression analysis was performed by aggregating cells belonging to the same sample and the same cluster to create a pseudo bulk sample. Then the negative binomial test implemented in R package DESeq2 was used to test the differential expression between groups of pseudo bulk samples. A subset of the scRNA data consisting of the epithelial cells was further analyzed by R studio Monocle3 package to determine a cell lineage trajectory^28^. The dimension reduction and UMAP coordinates performed using the Seurat package were transferred at the same time. The epithelial cells were clustered and partitioned using the Leiden algorithm for community detection^29^. Cluster 5 was set as the root and the single-cell trajectory was then built to show the transitions of cell transcriptomes.

### Cluster Identification and Cell Prediction

Differentially expressed genes for each cluster were obtained from the Seurat output which included average expression level (avg_log2FC), the fraction of cells within a cluster expressing a gene (pct.1), the fractions of genes outside of the cluster expressing a gene (pct.2), and the adjusted p-value. To capture marker genes per cluster, gene lists were produced after filtering for: minimum gene expression within a cluster (pct.1 > 1), cluster enrichment (pct.1-pct.2 > 0.15), effect size (avg_log2FC > 0.25) and statistical significance (p_val_adj <0.05). To reduce confounding signal, genes associated with the ribosome (*RPL*, *RPS*), mitochondria (*MT*-) or cell-cycle (*MKI67, TOP2A, CDK1, PCNA*) were excluded. Two gene lists per cluster were produced which included the top 50 genes ranked by expression magnitude (avg_log2FC) or cluster specificity (pct.1-pct.2). Next the marker gene lists were analyzed using Enrichr (Supp. Table 1)^30^. The top 2 terms, based on p-value, from the following databases were listed per cluster: CellMarker 2024, Tabula Sapiens, Human Gene Atlas, GO (Gene Ontology) Biological Process 2025, Reactome Pathways 2024, and KEGG 2026. To determine the degree of agreement between highly expressed genes and highly specific genes (genes expressed in the greatest number of cells per cluster), spearman rank correlation was calculated between both gene lists. Values ranked between 0-1 with values closer to 1 signifying higher agreement. Finally, the gene lists were combined with duplicates removed to create an overall marker list and entered into GPT4 for unbiased cluster prediction^31^. For epithelial clusters, marker genes were validated by immunohistochemistry in intact, premenopausal human vaginal as described above.

### Statistical Analysis

Graphpad Prism 10 was used for statistical analysis and graphing. Data with at least 3 biological replicates was tested for normality with Shapiro-Wilk test and when normal, Student’s t-test or multiple-comparison ANOVA was applied when appropriate. Non-parametric data was analyzed with Mann-Whitney tests or Kruskal-Wallis ANOVA. For all statistical analyses, a p-value of <0.05 was considered significant.

## Results

### The premenopausal vaginal epithelium contains epithelial cells with self-renewal and differentiation potential

To determine if the premenopausal human vagina contains epithelial cells with stem cell characteristics of self-renewal and differentiation, we repurposed clonogenic and organoid *in vitro* assays previously developed for endometrial stem cells^32,33^ (Fig. 1A, B, Table 1). From a heterogenous population of vaginal epithelial cells seeded and cultured for 15 days, we observed multiple, densely packed colonies formed with an efficiency of 4.04 ± 2.92% (n= 17 samples) (Fig. 1C). To determine if cells in the vaginal epithelium contained cells capable of differentiation, we seeded heterogenous single epithelial cells into Matrigel domes and cultured with differentiation media. We observed abundant organoids after 12 days with an efficiency of 7.67 ± 5.37% (n=13 samples) (Fig. 1D). Histologic analysis confirmed that organoids contained cells representing all layers of the intact, vaginal epithelium (Fig. 1E, F). The vagina contains a stratified squamous epithelium that has spatially distinct layers with basal cells giving rise to suprabasal cells - parabasal, intermediate, superficial - from basement membrane to lumen, respectively (Fig. 1C). We utilized TP63, a classic marker of progenitors in stratified squamous epithelial tissues^34,35^, and KRT5, a marker of basal and parabasal cells in human fetal vagina^22^ and skin^36^. In contrast to fetal vagina and skin, we observed KRT5 expression throughout the intact human vaginal tissues and organoids, whereas TP63 was restricted to basal and parabasal cells in the intact vagina and the periphery of the organoids (Fig. 1G, H). Similarly, we utilized E-cadherin, a component of adherens junctions that is present throughout the epithelium^37^ and strong in the basal and parabasal layers, and KI67, a classic marker of proliferative cells. In intact human vagina, we observed E-cadherin throughout the epithelium and KI67 in the parabasal cells (Fig. 1I). The organoids mimic the protein localization in intact tissues, where E-cadherin is present on the cell surface of all cells and KI67 localizes to the periphery of the organoid (Fig. 1J). Taken together, we illustrate the adult human vaginal epithelium contains cells with stem-like behavior as they exhibit self-renewal and differentiation potential in clonogenic and organoid assays, *in vitro*.

### Clonogenic and differentiation potentials reside specifically in basal vaginal epithelial cells marked by ITGA6 or NGFR

In stratified squamous epithelia, the basal cells contain the progenitor population responsible for tissue homeostasis and regeneration^38^. We isolated and fractionated vaginal basal epithelial cells by fluorescent-activated cell sorting (FACS) with CD9, a cell surface marker widely expressed in epithelia^39^, including endometrium^40^; and ITGA6 (CD49f), a cell-surface adhesion receptor that anchors basal cells to the basement membrane and is an established stem cell marker in many tissues^41^ (Fig. 2A). We captured 4 distinct cell populations CD9+ITGA6+ (16.7% of live cells), CD9+ITGA6- (7.96%), CD9-ITGA6+ (0.05%) and CD9-ITGA6- (74.8%) and determined that about 25% of cells express one or both markers (Fig. 2A, B). Our preliminary data indicates that colony forming activity and differentiation potential in organoids resides exclusively in the CD9+ fraction of the human vagina (Supp. Fig. 2). Next, we fractionated the CD9+ fraction based on expression of ITGA6. Colony forming activity was present exclusively in the CD9+ITAG6+ population with an efficiency of 4.15%, whereas colony forming activity was depleted in the CD9+ITGA6- (0.01%) (Fig. 2C, D). Next CD9+ITGA6+ and CD9+ITGA6- subpopulations were seeded in Matrigel and cultured with differentiation factors for 12 days. We observed more organoids formed in the CD9+ITGA6+ population with an efficiency of 2.13 % than the CD9+ITGA6- population (0.13%, Fig. 2E, F). These initial results illustrate that ITGA6 captures a subpopulation of CD9+ epithelial cells displaying a trend toward increased colony forming activity and organoid formation. Next, we tested NGFR (CD271), which marks a population of basal keratinocyte stem cells in the skin, lung, esophagus, oral mucosa, and is a stem cell marker in mouse vagina^42–46^. We used FACS to fractionate the CD9+ epithelial cells into four subpopulations based on NGFR expression levels: CD9+NGFR+ (12.64 ± 7.98%), CD9+NGFR- (24.35 ± 1.86%), CD9-NGFR+ (0.07 ± 0.04%), and CD9-NGFR- (63.0 ± 8.30%) and determined that roughly 37% of cells express one or both markers (Fig. 2G and H). Colony forming potential was greater in CD9+NGFR+ population with an efficiency of 7.21 ± 3.72% compared to 0.54 ± 0.58% in the CD9+NGFR- population (Fig. 2I, J, p<0.05). However, both NGFR+ and NGFR- subpopulations of CD9+ cells formed organoids after 12 days with equivalent efficiency, 7.00 ± 1.22 and 6.22 ± 3.07%, respectively (Fig. 2K and L, p>0.05). Taken together, we found both ITGA6+ and NGFR+ fractions of CD9+ epithelial cells have colony forming activity, whereas organoid forming activity was found in the ITGA6+, NGFR+, and NGFR- populations of CD9+ epithelial cells, in vitro.

**Figure 2.**
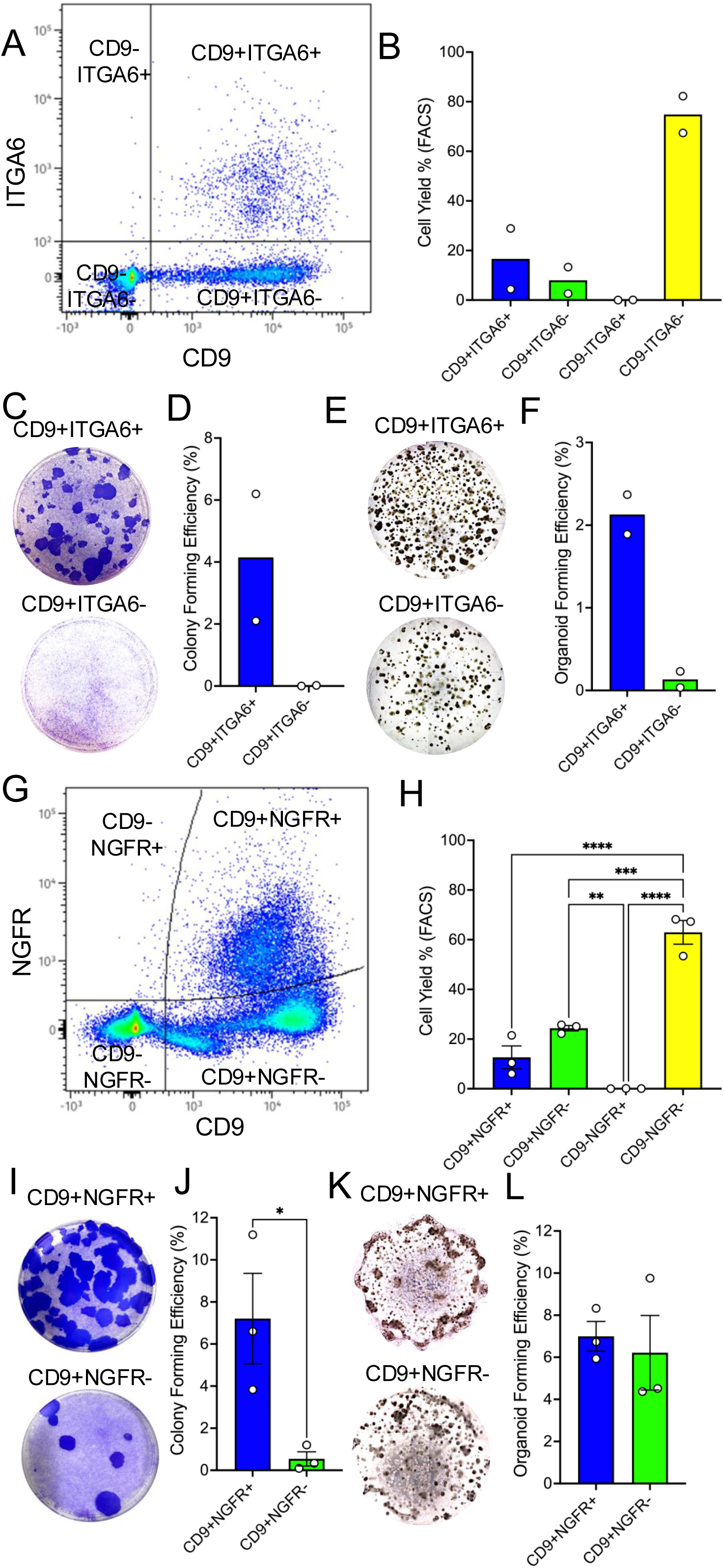
Stem cell markers, ITGA6 and NGFR, capture epithelial cells from the premenopausal human vagina capable of colony and organoid formation *in vitro*. A) Representative FACS plot displaying cells sorted by CD9 (broad epithelial cell marker) and ITGA6 (putative stem cell marker) from the vaginal epithelium. B) Cell yields (%) after FACS with ITGA6 and CD9 in the epithelium of the premenopausal vagina (n=2 samples). C) Representative culture plates stained with Crystal Violet seeded with single cells (1500/plate) from the CD9+ITGA6+ and CD9+ITGA6- subpopulations from the premenopausal vaginal epithelium and cultured for 14 days. D) Within CD9+ epithelial cells, ITGA6+ cells display colony forming efficiency (%) compared to ITGA6- cells (n=2 samples). Samples were triplicates and the average per sample is plotted. Statistical analysis not performed due to n-value. E) Representative images of Matrigel domes containing veOrganoids formed from CD9+ITGA6+ and CD9+ITGA6-subpopulations of vaginal epithelial cells. Imaged by transmitted light using a 4X objective. F) Organoid forming efficiency (%) of ITGA6+ and ITGA6- subpopulations of CD9+ vaginal epithelial cells (n=2). Samples were run in triplicate and the average per sample is plotted. Statistical analysis was not performed due to n-value. G) Representative FACS plot displaying subpopulations of vaginal epithelial cells using CD9 (broad epithelial cell marker) and NGFR (putative stem cell marker). H) Cell yields (% ± SEM) of epithelial subpopulations captured after cell sorting with NGFR and CD9 (n=3 samples, Kruskal-Wallis ANOVA, p<0.05). I) Representative colonies formed after 14 days in culture after CD9+NGFR+ or CD9+NGFR-single cells were plated (1500 cells/plate). J) CD9+NGFR+epithelial cells displayed increased colony forming efficiency compared to CD9+NGFR- cells after 14 days (n=3 samples, unpaired t-test, p<0.05). Samples were triplicates and the average plotted. K) Representative images of Matrigel domes containing organoids formed after 12 days from CD9+NGFR+ and CD9+NGFR-epithelial cells. Imaged by transmitted light using a 4X objective. L) CD9+NGFR+ and CD9+NGFR- epithelial cell populations showed similar organoid forming efficiency (% ± SEM, unpaired t-test, p>0.05).

### A transcriptomic atlas of the premenopausal human vagina

To date there is no transcriptomic atlas of human, adult, premenopausal, non-prolapsed vaginal tissue. To discover, validate and characterize markers of the stem/progenitor compartment and differentiated epithelial cell types that make up the human vagina, we performed single-cell RNA sequencing of full-thickness vaginal biopsies from premenopausal (median age 36 ± 7.0 years, N=6) women (Supp. Fig. 2). We profiled a total of 51,193 cells with 5717 unique Gene IDs and identified 26 transcriptionally distinct cell clusters (Fig. 3A, Supp. Table 1). Using the top 50 genes based on expression level and cluster specificity, we identified 4 major cluster types in the human vagina: epithelial (Fig. 3B), fibromuscular (Fig. 3C), endothelial (Fig. 3D), and immune (Fig. 3E)(Supp. Table 1). We focused further experiments specifically on the vaginal epithelium (Fig. 3B).

**Figure 3.**
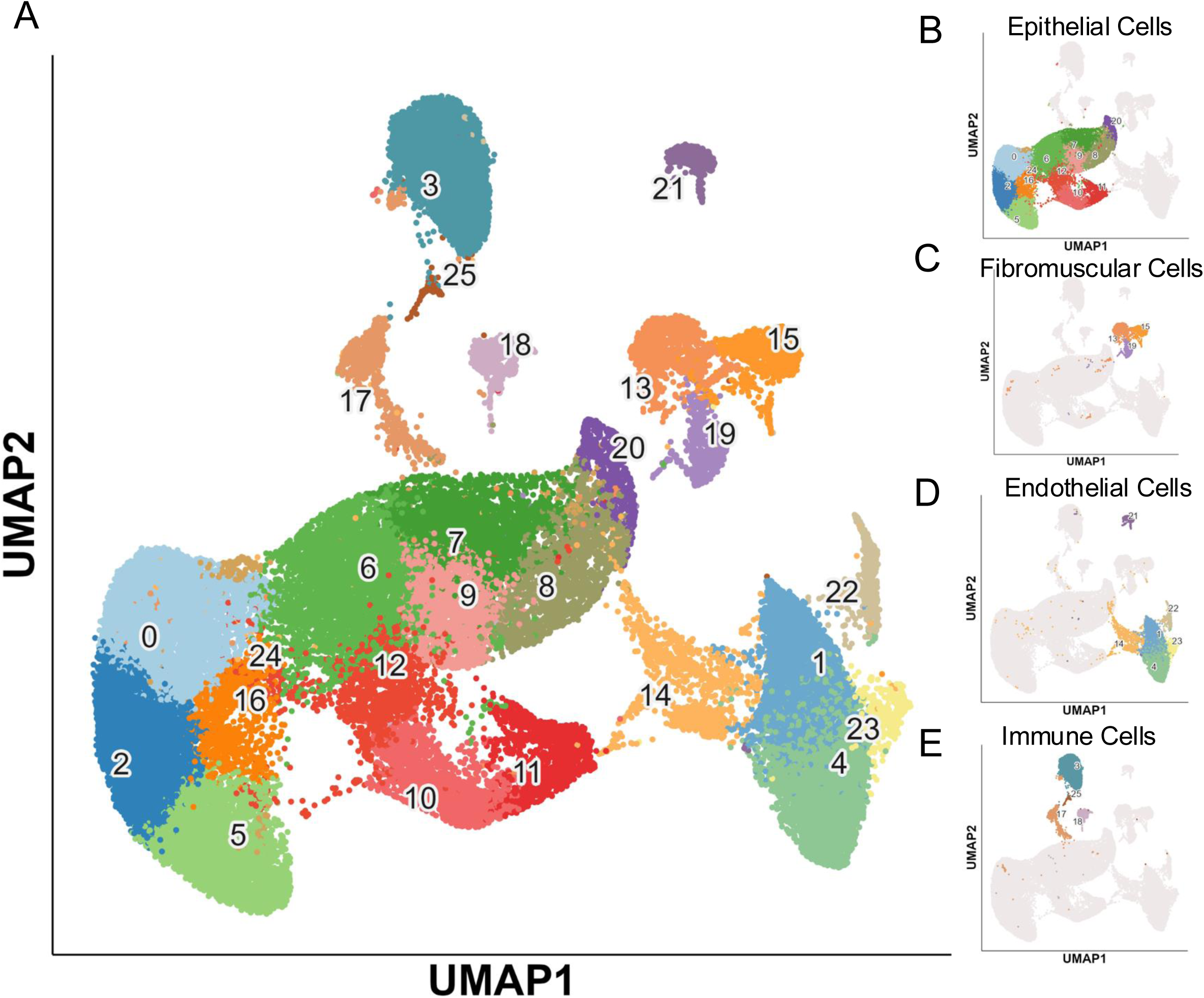
Single cell transcriptomic atlas of the premenopausal, non-prolapse, human vagina. A) UMAP displaying 26 cell clusters derived from single cell RNA sequencing (51,882 cells) of the human vagina (n=6 samples, age 25-43 years). B) UMAP of epithelial cell clusters (0, 2, 5, 6, 7, 8, 9, 10, 11, 12, 16, 20, 24). C) UMAP of fibromuscular cell clusters (13, 15, 19). D) UMAP of endothelial cell clusters (1, 4, 14, 21, 22, 23). E) UMAP of immune cell clusters (3, 17, 18, 25).

### Epithelial lineage trajectory and differentiation in the human vagina

The vaginal epithelium is stratified into multiple layers; however, the cell identities and state transitions are not well-defined. Using hierarchical clustering of the most prevalent genes within the predicted epithelial clusters, we identified 3 major histologic compartments which include basal, parabasal, and intermediate/superficial epithelial cells (Fig. 4A). Basal clusters (0, 2, 5, 16 and 24) display prevalence and expression of canonical basal epithelial marker, *COL17A1*^22^, parabasal clusters (6, 10, 11, 12) display reduced basal gene expression with induction of canonical differentiation genes such as *CLCA2*^47^ and *DSG1*^48^, and characterized by high expression of proliferation markers such as *TK1*^49^*, PCLAF*^50^, and *BIRC5*^51^. Clusters 7, 8, 9, 20 contain intermediate and superficial epithelial cells as they lack basal and proliferative cell markers and contain classic terminal differentiation markers such as *IVL*^52^ and *CRNN*^53^. Our preliminary data with NGFR+ epithelial cells from the human vagina (Fig. 2) as well as published reports on the mouse vagina suggest that NGFR marks a population of stem/progenitor cells in the human vaginal epithelium^21^. We performed pseudotime analysis using Monocle3 on the epithelial clusters (n=30,520 cells), which infers a biological progression and predicts lineage branching (Fig. 4B). Cluster 5 contains the most cells expressing *NGFR* (29.2%) relative to all other profiled cells in the vagina (3.6%) and compared to clusters that share basal epithelial cell signatures, including cluster 0 (6.08%), 2 (13.4%), 16 (12.1%), 24 (15.7%). Therefore, we set cluster 5 as the root for trajectory analysis (Fig. 4B). We observed trajectory loops within clusters 0, 2, 5, branching from 16 and 24 into clusters 6, 7, 8, 9, 10, 11, 12, 20 which suggests cells transition among basal cell clusters followed by a linear differentiation pathway (Fig. 4C and D). We predict NGFR marks a subpopulation of basal cells with stem/progenitor activity (clusters 0, 2, 5) that undergo initial differentiation (clusters 16 and 24) and transit-amplification (clusters 6, 10, 11, 12) within the parabasal compartment and fully differentiate into intermediate (clusters 7, 8, 9) and terminally differentiated superficial epithelial cells (cluster 20, Fig. 4C and D).

**Figure 4.**
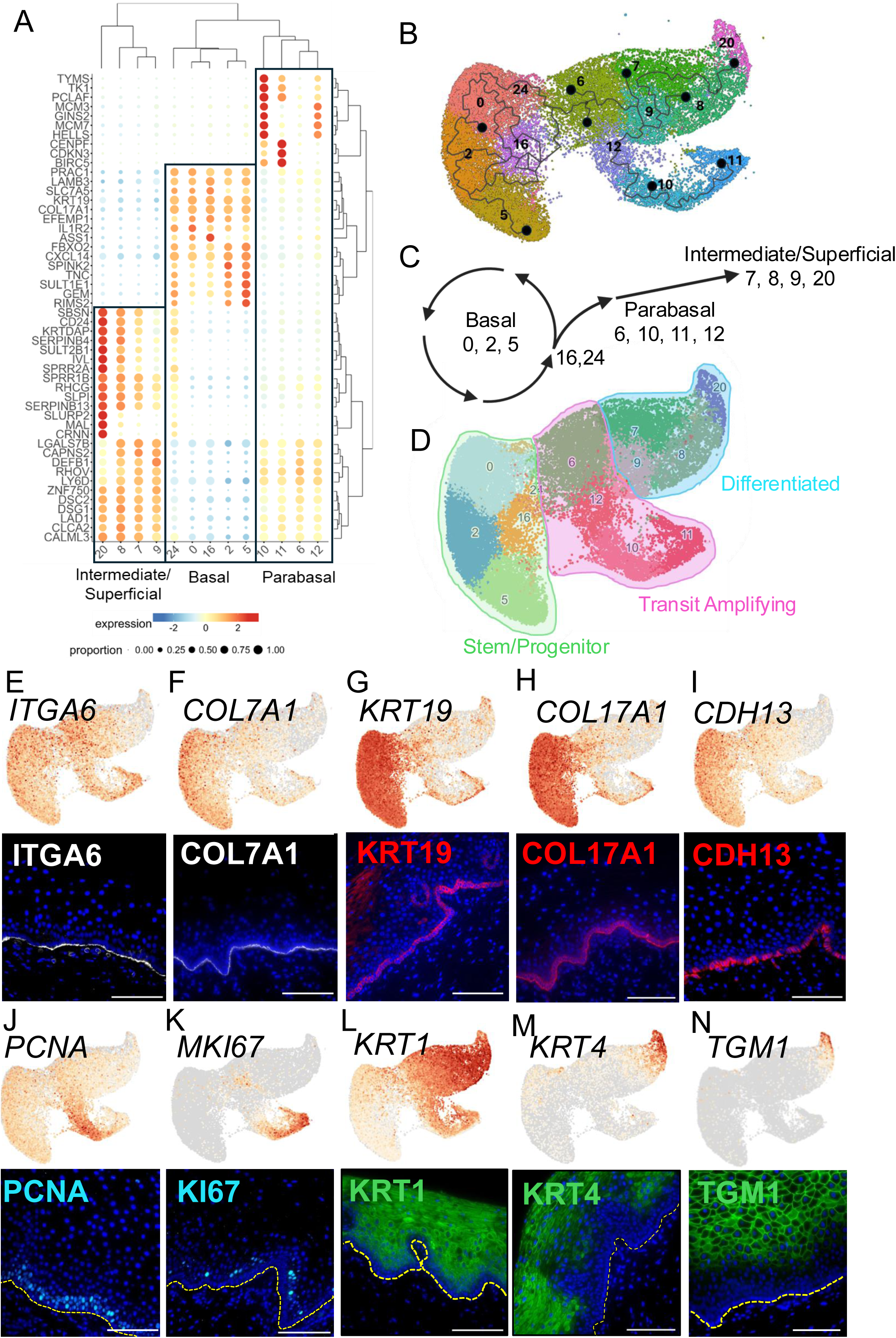
Characterizing the human premenopausal vaginal epithelium. A) Dotplot heatmap of most prevalent (pct.1-pct.2) genes per epithelial cluster results in hierarchal clustering of 3 major epithelial cell types (Basal, Parabasal, Intermediate/Superficial). B) Cell trajectory analysis of epithelial cell clusters (n=30,550 cells) after Monocle3 analysis. Cluster 5 was set as the root. C) Hypothesized differentiation pathway of epithelial cells within the vaginal epithelium. D) Hypothesized cell state overlayed on vaginal epithelial UMAP. E) UMAP of *ITGA6* gene expression (51.5%) in epithelial clusters and basement membrane protein localization of ITGA6 (white) in the premenopausal vaginal epithelium. F) UMAP of *COL7A1* gene expression (37.4%) and basement membrane protein localization of COL7A1 (white) in the premenopausal vaginal epithelium. G) UMAP of *KRT19* gene expression (68.6%) and basal epithelial cell protein localization of KRT19 (red) in the premenopausal vaginal epithelium. H) UMAP of *COL17A1* gene expression (65.7%) and basal epithelial cell protein localization of COL17A1 (red) in the premenopausal vaginal epithelium. I) UMAP of *CDH13* gene expression (51.0%) and basal epithelial cell protein localization of CDH13 (red) in the premenopausal vaginal epithelium. J) UMAP of *PCNA* gene expression (49.9%) and basal/parabasal epithelial cell protein localization of PCNA (cyan) in the premenopausal vaginal epithelium. K) UMAP of *MKI67* gene expression (10.7%) and parabasal epithelial cell protein localization of KI67 (cyan) in the premenopausal vaginal epithelium. L) UMAP of *KRT1* gene expression (72.8%) and suprabasal epithelial cell protein localization of KRT1 (green) in the premenopausal vaginal epithelium. M) UMAP of *KRT4* gene expression (15.7%) and intermediate/superficial epithelial cell protein localization of KRT4 (green) in the premenopausal vaginal epithelium. N) UMAP of *TGM1* gene expression (3.89%) and superficial epithelial cell protein localization of TGM1 (green) in the premenopausal vaginal epithelium. cells. Scale bars are 100 μm.

We used immunohistochemistry to validate the single cell transcriptional data and identify markers of the different compartments of the human vaginal epithelium: basal cells, proliferative cells, and suprabasal cells which include parabasal, intermediate and superficial epithelial cells (see Fig. 1E for reference). We selected markers from the literature as well as markers with high specificity (pct. 1 values, proportion expressing cells within a cluster) or enrichment (logFoldChange, gene expression level in cluster) in our scRNAseq data and performed cross-platform immunohistochemical validation with intact, premenopausal tissue. Both ITGA6 and COL7A1 mark the basement membrane of the vaginal epithelium in humans (Fig. 4E, F). KRT19, COL17A1, CDH13 mark cells along the basement membrane with gene expression specificity to clusters 0 (97.4, 95.8, 52.3%), 2 (98.7, 98.2, 63.3%), 5 (99.5, 99.9, 72.6%), 16 (98.4, 99.7, 78.0%) and 24 (97.7, 94.1, 55.3%), respectively (Fig. 4G, H, I). Next, we analyzed proliferation markers, PCNA and KI67. We observed a broader expression of *PCNA* (18.2-80.4% of cells expressing across all clusters) compared with *MKI67*, which was relatively restricted to the parabasal region (clusters 6, 9.78%; 9 3.43%; 10, 70.0%; 11, 91.8%; 12, 11.2%) of the vaginal epithelium. Protein expression for both markers was most prominent in the parabasal compartment (Fig. 4J, K), supporting the assignment of these clusters to the transit amplifying compartment of the human vaginal epithelium (Fig. 4D). Next, we investigated differentiation markers, KRT1, KRT4 and TGM1. KRT1 shows suprabasal protein localization, predominantly marking the parabasal (cluster 6, 92.4%; 10, 89.1%; 11, 90.7%; 12, 88.2%), intermediate (cluster 7, 99.8%; 8, 100.0%; 9 (99.7%)) and superficial (cluster 20, 98.0%) epithelial cells whereas KRT4 is specific to intermediate (clusters 7, 28.4%; 8, 54.2%) and superficial cells (cluster 20, 92.9%). TGM1 is restricted to superficial cells (cluster 20, 73.8% expression) (Fig. 4L, M, N). Together we illustrate cluster validation of the vaginal epithelium informed by the single cell transcriptomic analysis and present a catalog of markers to define the basement membrane, basal and suprabasal cells, which include proliferative, parabasal, intermediate and superficial epithelial cells.

### NGFR+ITGA6+ epithelial cells from the premenopausal vagina display exhibit self-renewal in vitro

Our initial in vitro assays demonstrate that *in vitro* stem cell activity resides in the ITGA6+ and NGFR+ fractions of the human vagina. However, our scRNAseq data indicate that NGFR+ cells are restricted to the basal compartment and represent a smaller subpopulation of ITGA6+ cells which are present in the basal and parabasal compartments (Fig. 5A). Over half (51.5%) of all epithelial cells in the human vagina express *ITGA6* whereas a smaller fraction express *NGFR* (8.9%) (Fig. 5B and C). To ensure capture of an epithelial cell population, we utilized CD9 as a broad marker expressed in 99.0% of the cells within the epithelial clusters, confirmed by epithelial cell specific protein localization (Fig. 5C) and preliminary data suggesting CD9- populations lack stem cell activity (Supp. Fig. 2). We then used FACS and *in vitro* stem cell assays to test the hypothesis that stem cell activity resides in the NGFR+ fraction of CD9+ITGA6+ cells in the human vagina. CD9+ epithelial cells (Fig. 5D, E) were fractionated based on ITGA6 and NGFR expression (Fig. 5 F). The majority of CD9+ epithelial cells were ITGA6+NGFR+ (61.1 ± 16.3%) with lower percentages in the ITGA6+NGFR- (15.2 ± 5.16%) and ITGA6-NGFR- (26.2 ± 13. 4%) fractions (Fig. 5F, G). These fractions were tested in the *in vitro* stem cell assays. There were almost no cells in the ITGA6-NGFR+ (0.33 ± 0.17%) fraction, indicating that all NGFR+ cells are in the ITGA6+ fraction, so this fraction was not tested (Fig. 5F, G, p<0.05). We found that the ITGA6+NGFR+ cells had significantly higher colony forming efficiency (3.32 ± 2.43%) after 14 days in culture than the ITGA6+NGFR- (0.26 ± 0.25%) and ITGA6-NGFR- (0.006 ± 0.02%) cells (Fig. 5H, I, p<0.05).

**Figure 5.**
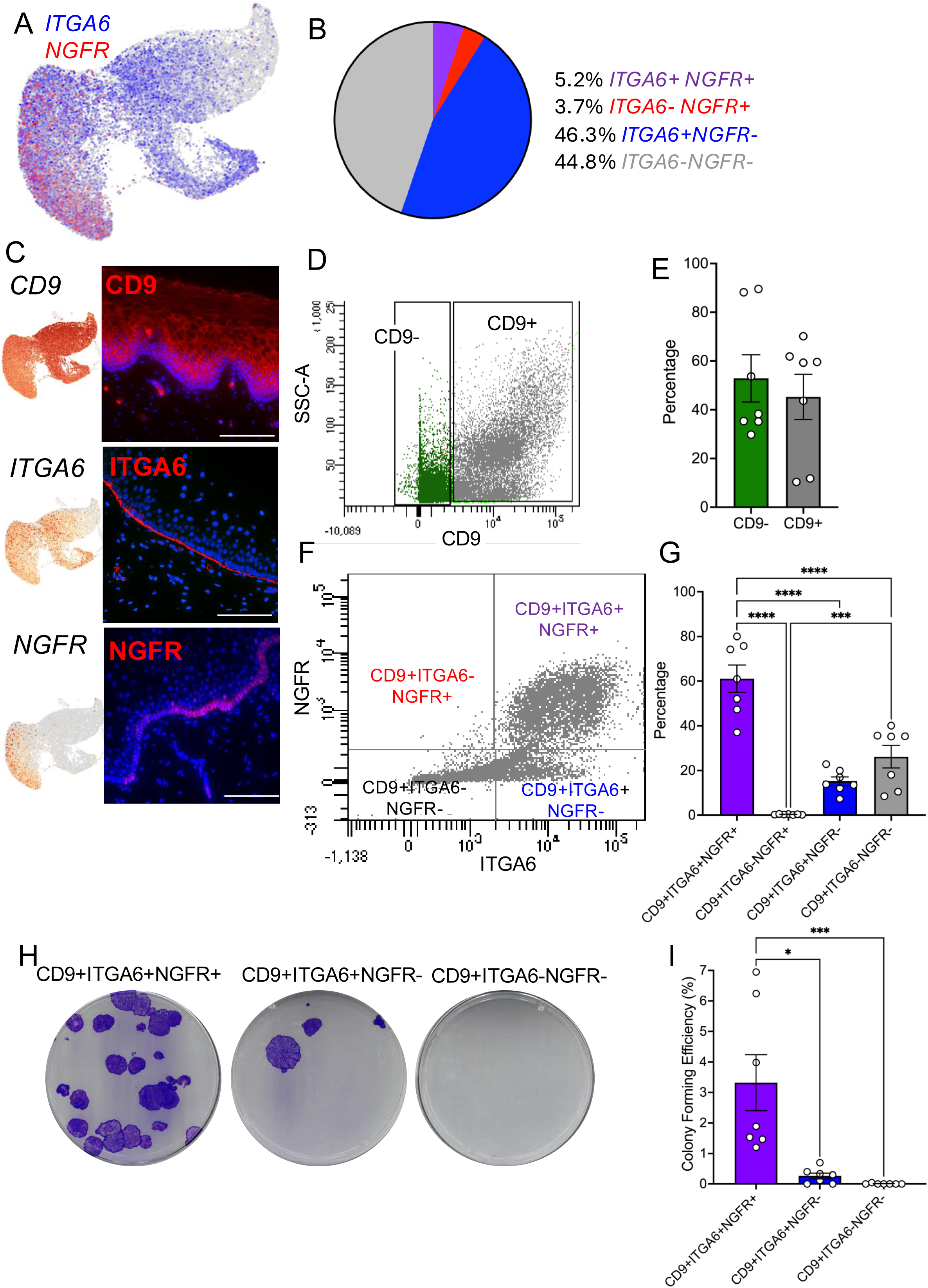
NGFR marks a subpopulation of ITGA6+ basal epithelial cells in the premenopausal vagina that has enhanced colony forming potential. A) Co-expression UMAP of epithelial clusters illustrates subpopulation of NGFR (red) cells and ITGA6 (blue) cells. B) Quantification of epithelial cell expression of NGFR and ITGA6 in the vaginal epithelial clusters. C) Gene expression of CD9 (99.0%), ITGA6 (51.5%) and NGFR (8.91%) in epithelial clusters and matched immunohistochemistry (red) revealing protein expression in the vaginal epithelium. D) Representative FACS plot of CD9 subpopulations. E) Quantification (% ± SEM) of cell yields when sorted for CD9+ epithelial cells (n=7 samples, unpaired t-test, p>0.05). F) Representative FACS plot of NGFR and ITGA6 subpopulations within CD9+ epithelial cells. G) Cell yields (% ± SEM) of NGFR and ITGA6 subpopulations within CD9+ cells (n=7 samples, Kruskal-Wallis ANOVA, p<0.05). H) Representative clonogenicity plates of CD9+ITGA6+NGFR+, CD9+ITGA6+NGFR- and CD9+ITGA6-NGFR- subpopulations after 14 days in culture. I) CD9+ITGA6+NGFR+ epithelial cells are the most efficient at colony formation (% ± SEM) compared to CD9+ITGA6+NGFR- and CD9+ITGA6-NGFR- (n=7 samples, Kruskal-Wallis, ANOVA, p<0.05).

### NGFR+ progenitor epithelial cells give rise to fully differentiated vaginal epithelial cells in organoids, in vitro

We next tested whether NGFR+ cells could establish organoids and produce cells representing all compartments of the vaginal epithelium. After FACS, we seeded CD9+ITGA6+NGFR+, CD9+ITGA6+NGFR-, CD9+ITGA6-NGFR- cells into Matrigel and cultured with differentiation factors for 10-12 days. Organoid forming efficiency was higher in the CD9+ITGA6+NGFR+ subpopulation (2.58 ± 1.76%) than in the CD9+ITGA6+NGFR- fraction (0.44 ± 0.24%) and CD9+ITGA6-NGFR- (0.06 ± 0.09%) fraction of CD9+ epithelial cells derived from premenopausal human vagina (Fig. 6A, B, p<0.05). Vaginal epithelial organoids (veOrganoids) have biologic similarity to intact premenopausal tissue. We observed conservation of ITGA6 and CDH13 representing basal cell populations around the exterior of the organoids as well as KI67+ cells indicating maintenance of proliferation (Fig. 6C). Furthermore, we observed stratified epithelial cell organization within veOrganoids as both basal, stained by CDH13, and differentiated TGM1 positive epithelial cells are present with exclusive protein localization in the exterior and interior compartments, respectively (Fig. 6D). In addition to quantitative differences in the number of organoids, we observed a difference in CD9+ITGA6+NGFR+ organoid diameter (191.7 ± 42.2 μm) compared to CD9+ITGA6+NGFR-organoids (161.1 ± 37.9 μm)(Fig. 6E, F, p<0.05). We observed no obvious morphologic differences between NGFR+ and NGFR- organoids (Fig. 6G). Next, we tested whether NGFR+ and/or NGFR- cell-derived organoids contained all cell types of the human vaginal epithelium. Both NGFR+ and NGFR- organoids contain CD9 expression in all cells and ITGA6+ cells around the periphery of the organoid (Fig. 6H, white). In addition, NGFR+ and NGFR- organoids contained cells expressing the basal cell markers, COL17A1 and CDH13, around the periphery of the organoid (Fig. 6H, red). However, NGFR+ derived organoids contained both KRT4+ and TGM1+ cells in the interior of the organoid demonstrating differentiation to the intermediate and superficial epithelial cells; whereas NGFR- derived organoids lack consistent KRT4 and TGM1 cells (Fig. 6H, green). Taken together, we demonstrated NGFR+ putative progenitor cells derived from premenopausal human vagina can fully differentiate in vitro while preserving a basement membrane, basal cells, and proliferation which supports recapitulation of the major cell compartments of the intact vaginal epithelium.

**Figure 6.**
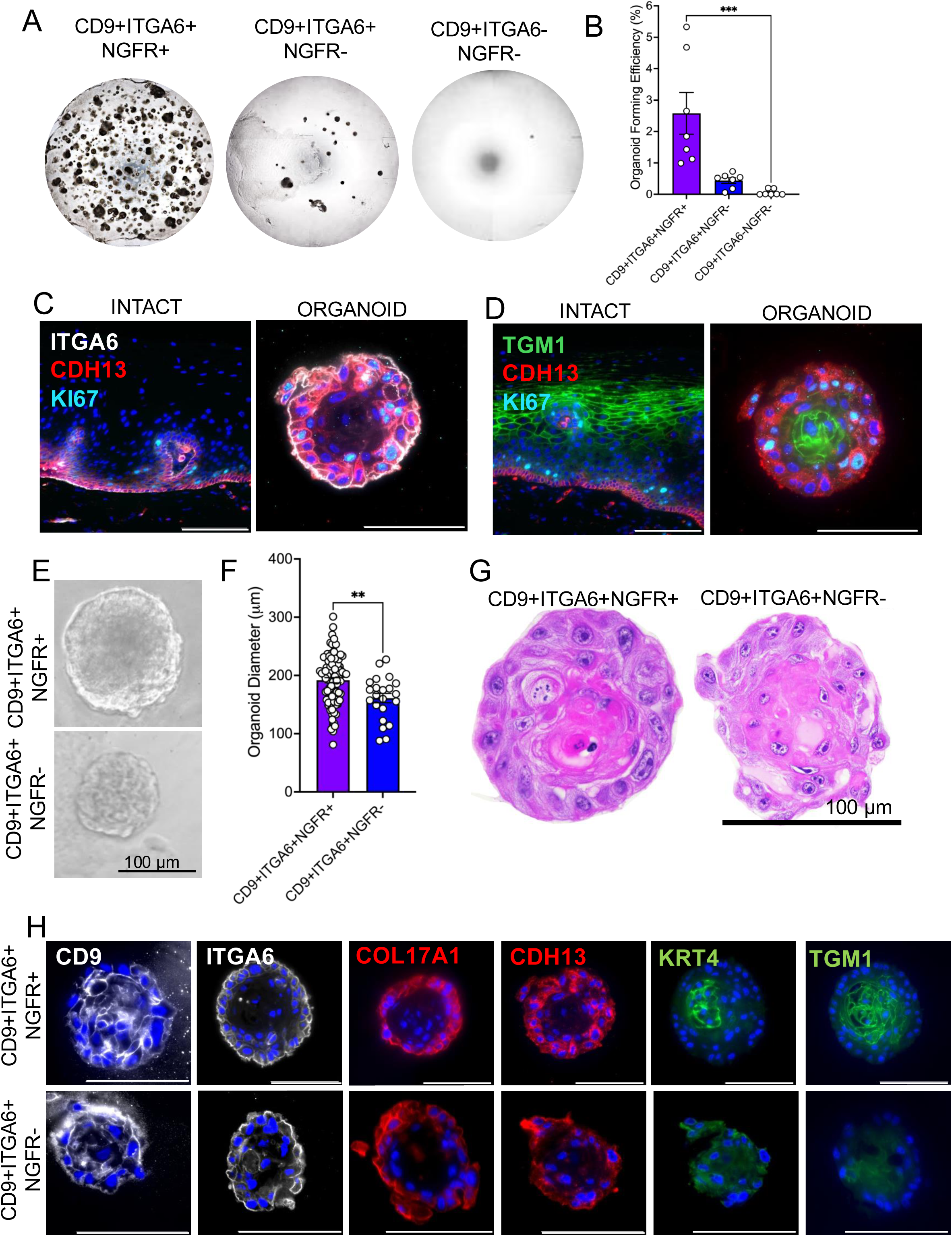
NGFR+ cells produce organoids that maintain basal cells and give rise to superficial epithelial cells. A) Representative images of Matrigel domes seeded with CD9+ITGA6+NGFR+, CD9+ITGA6+NGFR-, CD9+ITGA6-NGFR- progenitor cells (5000 cells/dome) and cultured for 10-12 days. Images with transmitted light with a 4X objective. B) CD9+ITGA6+NGFR+ progenitor subpopulation are more efficient at organoid formation (% ± SEM) in vitro compared to CD9+ITGA6+NGFR- and CD9+ITGA6-NGFR- subpopulations (n=7 samples, Kruskal-Wallis ANOVA, p<0.05). C) Representative image of intact vaginal epithelium and organoids co-stained with ITGA6 (white), CDH13 (red), and KI67 (cyan). Imaged with fluorescent microscopy and 40X objective. Scale bar, 100 µm. D) Representative images of intact vaginal epithelium and organoids co-stained with CDH13 (red), MK67 (cyan) and TGM1 (green). Imaged with fluorescent microscopy and 40X objective. Scale bar, 100 µm. E) Representative image of a single organoid cultured for 12 days imaged with transmitted light and a 40X objective. Scale bar is 100 µm. F) Diameter (μm ± SEM) of organoids derived from CD9+ITGA6+NGFR+ (n=92 organoids) and CD0+ITGA6+NGFR- (n=22 organoids) cell subpopulations, unpaired t-test, p<0.05). G) Representative images of cross sections of organoids stained with hematoxylin and eosin and imaged with brightfield and a 100X objective. H) Immunohistochemical validation of CD9 (white), ITGA6 (white), COL17A1 (red), CDH13 (red), KRT4 (green), TGM1(green) in organoids derived from CD9+ITGA6+NGFR+ and CD9+ITGA6+NGFR- progenitor cells. A single representative image is shown from a pool of 3-13 organoids analyzed per marker. Scale bar is 100 μm.

## Discussion

The intrinsic regeneration of the vaginal epithelium suggests the existence of epithelial stem cells which previously had not yet been identified in the human vagina. We validated the use of classic clonogenic^54^ and organoid^55^ assays for defining stem cell function for the human vagina. Using these assays that we developed for the endometrium^32,33^, we identified specific cellular populations in the human vaginal epithelium with the ability to self-renew and differentiate *in vitro*. Stem cell activity specifically segregated to cells on the basement membrane of the vaginal epithelium that express ITGA6 and NGFR, markers of stem cells in other tissues, including the mouse vaginal epithelium.

Our scRNAseq data of adult, premenopausal, non-prolapsed vagina provides the field a foundational dataset. We specifically defined the differentiation trajectory of the human vaginal epithelium and validated markers that span the full-thickness epithelium from basement membrane to superficial cells. We discovered that NGFR marks a subpopulation of cells on the basement membrane of the vaginal epithelium by both gene expression and protein validation and demonstrated that these cells are the likely stem/progenitors that self-renew and differentiate to maintain the vaginal epithelium. We identified CD9+ as a surface marker of vaginal epithelial cells isolated from premenopausal individuals and determined that NGFR and ITGA6 surface proteins segregated CD9+ cells into 4 subpopulations. The ITGA6+NGFR+ population was the most efficient at colony and organoid formation compared to ITGA6+NGFR- or ITGA6-NGFR- populations. While both ITGA6+NGFR+ and ITGA6+NGFR- populations generated organoids, only the ITGA6+NGFR+ organoids displayed terminal differentiation to KRT4 or TGM1 cells in the center while maintaining basal cells marked by COL17A1 and CDH13 at the periphery. These results strongly suggest that the human vaginal epithelium is a stem cell-based system and confirm that the NGFR+ subpopulation of basal cells in the human vaginal epithelium self-renewed and differentiated, properties of all adult tissue stem cells.

Our trajectory analysis of the premenopausal human vagina illustrates several trajectory loops within the basal cell clusters, suggesting the stem cell pool is heterogenous. Tissue-resident stem cells exist in specialized states such as quiescence, slow-cycling, primed or activated which are necessary to respond to environmental cues, ensuring tissue homeostasis and maintenance of tissue function over a lifetime^56^. We illustrate NGFR marks a subpopulation of basal cells capable of clonal expansion and differentiation; however, it is possible NGFR marks a particular state of veSC and future work is necessary to understand the heterogeneity of epithelial stem cells in the human vagina and the cues regulating transitions between stem cell states and differentiation to transit amplifying and terminally differentiated cells.

The vaginal epithelium is sensitive to endocrine changes across the menstrual cycle and the lifespan of women, and these contribute to an individual’s susceptibility to viral and bacterial infections as well as atrophy^57–59^. We achieved epithelial differentiation to superficial epithelial cells *in vitro* without the use of serum or steroid hormones including estrogen. While there is a need to develop non-hormonal therapies as less than 5% of menopausal women use estrogen therapy^60^, it is also necessary to determine how estrogen and sex steroids mediate veSC function particularly through developmental milestones such as menarche, childbirth and the menopausal transition.

Stem cells exist in specialized niches^61^. While the focus of our study was identification of epithelial stem/progenitor cells, our novel transcriptomic atlas of the full thickness premenopausal human vagina also identified cell clusters specific to fibromuscular, endothelial and immune cells. Vaginal dysbiosis, inflammation and fibrosis are features of the aging vagina and alterations in the underlying mesenchyme and the immune cell composition may contribute to atrophy and stem cell exhaustion in the epithelium^62–66^. Furthermore, understanding the environmental cues that emanate from these tissues may provide clues about paracrine mechanisms that regulate stem cell function or tissue regeneration in all compartments of the human vagina.

Our experiments have commenced the building of a toolbox for studying human vaginal epithelium, including validated markers of epithelial stem cells and their differentiated progeny as well as *in vitro* stem cell assays. Our study identified cell surface markers that isolate basal progenitor cells that allowed sorting of viable, potentially therapeutic cells without genetic manipulation. These tools to enrich the endogenous epithelial stem cell population may lead to future treatments to maintain or restore the vaginal epithelium in individuals born with congenital malformation of the vaginal canal^67^ or who have undergone surgical procedures that removed vaginal tissue^68,69^, or needing vaginal rejuvenation during aging, menopause, or following chemotherapy or radiation therapy^70^.

In summary, we demonstrate that the human vaginal epithelium contains cells with stem cell function. We identified a subpopulation of vaginal epithelial cells (NGFR+) that exhibit typical stem cell characteristics of self-renewal and differentiation, including the potential to produce terminally differentiated vaginal epithelial cells in organoids. In the process, we developed and validated several experimental tools that will support future studies of the human vaginal epithelium: 1) a novel single cell atlas of the human, premenopausal vagina, 2) molecular markers of distinct cell types in the vaginal epithelium from basement membrane to superficial cells, and 3) in vitro veSC assays. Our study advances the fundamental understanding of stem cells and their progeny in the human vagina, which may lead to novel treatments for vaginal dysfunction and injury.

## Author Contributions

J.L.B. is responsible for preparation of the manuscript including generating figures and writing the text as well curating transcriptomic data and bioinformatic analysis as well as performing FFPE immunohistochemistry on intact tissue and veOrganoids. K.E.S. is responsible for sample processing, data acquisition, in vitro stem cell assays, OCT immunohistochemistry on intact tissue and veOrganoids from Hudson Institute of Medical Research as well as contributing to the preparation of the manuscript. T.M.S. is responsible for tissue acquisition and sample processing at the University of Pittsburgh, performing in vitro assays and preparing the manuscript. T.C. is responsible for processing and visualization of the scRNAseq data, contributing to marker selection and performing cell trajectory analysis. A.R.C. is responsible for sample processing, data acquisition, in vitro stem cell assays and OCT immunohistochemistry on intact tissue and veOrganoids from Hudson Institute of Medical Research. S.P.B. is responsible for sample processing, data acquisition and establishing the single cell organoid culture method. S.G. contributed to study conception and design and generation of preliminary data. K.R. is responsible for sample processing and data acquisition. C.E.G. is the scientific lead of the Hudson Institute team and contributed to the study conception, design and execution as well as editing and approving the manuscript for submission. K.E.O and P.A.M. are the scientific leads at the University of Pittsburgh, they contributed to study conception, design and execution of the study, writing the manuscript, approving for submission and handling correspondence.

## Acknowledgements

The authors thank those at the University of Pittsburgh and Monash Health who donated tissue and made this work possible. We are grateful to Drs. Natalie Brzoza, April Dunmyre, Erin Rhinehart, Jennifer Makin, Kelly DiMattio, Anna Rosamilia at Monash Health, Cabrini Health and Waverley Private hospital and their clinical research staff for assistance in specimen collection. We thank Lindsey Baranski and Kim Jarret for leading patient recruitment, as well as Gabriella King for tissue acquisition, processing and preparation at the University of Pittsburgh. Work performed in the University of Pittsburgh Single Cell Core Facility (RRID:SCR_025110) and services and instruments used in this project were graciously supported, in part, by the University of Pittsburgh, the Office of the Senior Vice Chancellor for Health Sciences. We would like to thank the Advanced Genomics Core at the University of Pittsburgh for assistance in processing the single cell transcriptomic data of the premenopausal, human vagina. We thank Sarah Munyoki at the University of Pittsburgh for assistance in visualization of transcriptomic data. We thank the Histology and Microimaging Core at Magee-Womens Research Institute and Pitt Biospecimen Core at UPMC Shadyside and the Monash Histology Platform for assistance in processing, embedding and sectioning human tissue and cells used in this study. We would like to thank the Cell Cytometry and Sorting Core at Magee-Womens Research Institute and FlowCore at Monash University/Hudson Institute for assistance in cell sorting experiments. Experimental schematics were made using BioRender.

## Supplemental Information

**Supp. Fig. 1.**
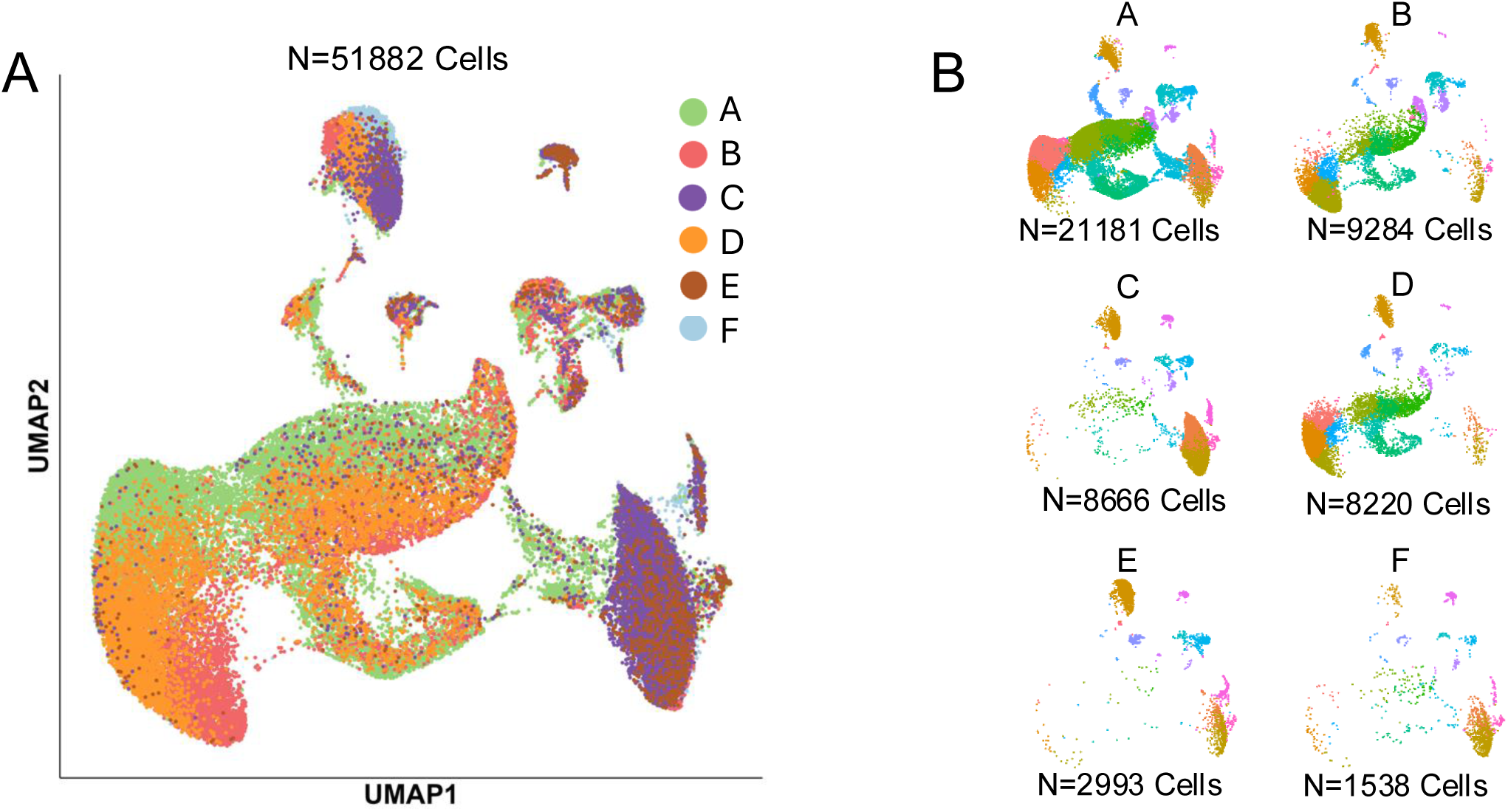
Sample specific contributions to transcriptomic map of the premenopausal vagina. A) UMAP displaying sample contribution to all clusters (n=6). B) Individual sample UMAPs display varying cell numbers and cluster contributions.

**Supp. Fig. 2.**
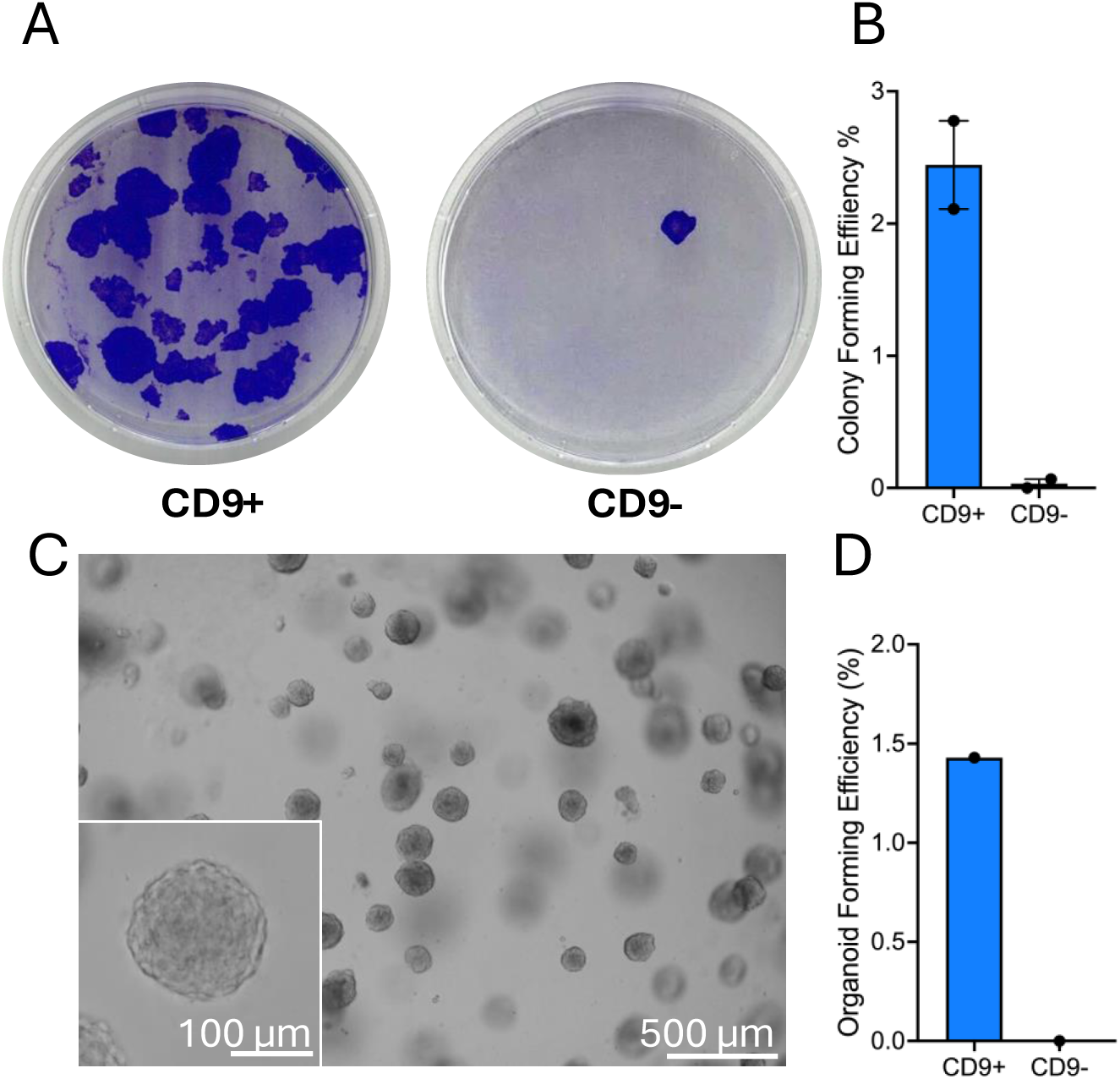
CD9+ epithelial cells contain clonogenic activity. A) Representative CFU plate of CD9+ITGA6+NGFR+ and CD9-ITGA6+NGFR+ cells after 12 days. B) Quantification of clonogenic activity from FACS sorted CD9+ and CD9- cells in the human vaginal epithelium (n=2). C) Representative brightfield image of CD9+ organoids. D) CD9+ epithelial cells give rise to organoids whereas CD9- cells are unable to form organoids (n=1).

<u>Supp. Table 1. Premenopausal vaginal cluster identification and prediction via EnrichR and GPT4.</u> Table contains combined top 50 marker genes by expression level (avg_log2FC) and cluster specificity (pct.1-pct.2) per cluster and corresponding terms from CellMarker 2024, Tabula Sapiens, Human Gene Atlas, GO Biological Process 2025, Reactome Pathways 2024 and KEGG 2026. Spearman correlation was performed across both gene lists to calculate concordance between highly expressed and enriched genes. Based on the combined lists, blinded prediction of cluster identity was performed via GPT4.

